# Gain without pain: Adaptation and increased virulence of Zika virus in vertebrate host without fitness cost in mosquito vector

**DOI:** 10.1101/2023.03.20.533515

**Authors:** Anna S. Jaeger, Jeffrey Marano, Kasen Riemersma, David Castañeda, Elise Pritchard, Julia Pritchard, Ellie K. Bohm, John J. Baczenas, Shelby L. O’Connor, James Weger-Lucarelli, Thomas C. Friedrich, Matthew T. Aliota

## Abstract

Zika virus (ZIKV) is now in a post-pandemic period, for which the potential for re-emergence and future spread is unknown. Adding to this uncertainty is the unique capacity of ZIKV to directly transmit between humans via sexual transmission. Recently, we demonstrated that direct transmission of ZIKV between vertebrate hosts leads to rapid adaptation resulting in enhanced virulence in mice and the emergence of three amino acid substitutions (NS2A-A117V, NS2A- A117T, and NS4A-E19G) shared among all vertebrate-passaged lineages. Here, we further characterized these host-adapted viruses and found that vertebrate-passaged viruses also have enhanced transmission potential in mosquitoes. To understand the contribution of genetic changes to the enhanced virulence and transmission phenotype, we engineered these amino acid substitutions, singly and in combination, into a ZIKV infectious clone. We found that NS4A- E19G contributed to the enhanced virulence and mortality phenotype in mice. Further analyses revealed that NS4A-E19G results in increased neurotropism and distinct innate immune signaling patterns in the brain. None of the substitutions contributed to changes in transmission potential in mosquitoes. Together, these findings suggest that direct transmission chains could enable the emergence of more virulent ZIKV strains without compromising mosquito transmission capacity, although the underlying genetics of these adaptations are complex.

## Introduction

The adaptive potential of RNA viruses is driven by error-prone replication which creates a heterogeneous swarm of virus genotypes within each host (1). RNA arthropod-borne viruses (arboviruses) therefore have the potential for rapid evolution but exhibit high degrees of consensus genome sequence stability in nature (summarized in (2)). The need to navigate distinct host environments and barriers to infection and transmission is believed to restrict arbovirus evolution. Indeed, the fitness trade-off hypothesis for arboviruses posits that fitness gains in one host come at the cost of fitness losses in the other host. Paradoxically, a preponderance of experimental evidence suggests that host alternation does not automatically limit the adaptive potential of arboviruses (reviewed in (3) and (4)). Rather, arboviruses may be the ultimate generalists, evolving mechanisms to limit or avoid the effects of trade-offs from dual-host cycling. They may be positioned to evolve rapidly when novel conditions arise.

This is perhaps best exemplified by the fact that many arbovirus epidemics have been associated with virus genetic change. For example, Indian-Ocean lineage chikungunya virus (CHIKV) adaptation to *Aedes albopictus* was mediated by a series of envelope glycoprotein E2/E3 and E1 substitutions during an explosive outbreak on La Reunion Island (5, 6). Other examples of epidemic-enhancing mutations include West Nile virus adaptation for more efficient transmission by North American mosquitoes (7) and Venezuelan equine encephalitis virus adaptations that produce high viral titers in horses (8). Finally, considerable research effort has been dedicated to identifying mutations that may have contributed to Zika virus (ZIKV) epidemic potential or contributed to the appearance of previously unobserved severe clinical manifestations (reviewed in (9)). Whether these mutations are important determinants of ZIKV emergence and spread remains an open question (10–12).

However, ZIKV is unique amongst mosquito-borne viruses due to its capacity to directly transmit between humans via sexual transmission. While it is epidemiologically challenging to quantify the contribution of sexual transmission to infection incidence in endemic regions, recent studies have shown that sexual transmission may be a more important driver of ZIKV transmission than previously thought (13–15). The ability to forgo mosquito-borne transmission for sexual transmission therefore creates a scenario where a highly mutable virus may be able to explore broader sequence space leading to the potential for novel host-adaptive mutations becoming fixed in the human population. To this end, we recently explored the consequences of releasing ZIKV from alternate host cycling between mosquito and vertebrate hosts. We modeled single and alternate host transmission chains in *Ifnar1^-/-^* mice and *Aedes aegypti* mosquitoes (16). We found that mouse-and mosquito-adapted lineages replicated to significantly higher titers in mice compared to unpassaged ZIKV, and that mouse-adapted viruses were universally lethal. Mosquito-adapted and alternate-passaged viruses had more heterogeneous phenotypes, with some lineages producing enhanced mortality rates relative to unpassaged virus, and other lineages resulting in little to no mortality (16). We also found that during direct vertebrate transmission chains, ZIKV consistently evolved enhanced virulence coincident with the selection of two polymorphisms at a single loci in NS2A—A117V and A117T—and at NS4A E19G and NS2A I139T (16). While these data demonstrate that ZIKVs experimentally evolved through direct vertebrate transmission chains possess increased virulence, we did not assess the potential trade-offs between virulence and transmission, nor did we functionally characterize these amino acid substitutions using reverse genetics.

To evaluate the extent to which experimental evolution of ZIKV through mice, mosquitoes, and alternate host passaging alters the vector competence phenotype in mosquitoes, we exposed *Ae. aegypti* to ZIKV from each passage series (n=2 lineages per passage series) and assessed infection, dissemination, and transmission potential and compared these rates to unpassaged ZIKV. We found that the mouse-passaged lineages had enhanced transmission potential in these mosquitoes compared to unpassaged, mosquito-passaged, or alternately-passaged ZIKV lineages. To investigate the viral genetic determinants for enhanced virulence and transmission for these ZIKV variants, we engineered the NS2A A117V, A117T, and NS4A E19G substitutions—singly and in combination—into the Puerto Rican ZIKV isolate PRVABC59 (ZIKV-A117V, ZIKV-A117T, ZIKV-E19G, ZIKV-A117V/E19G, and ZIKV-A117T/E19G) and infected mice and mosquitoes with these viruses. This is the same parental isolate on which these mutations emerged (16). Viruses containing NS4A-E19G had increased mortality and viremia, increased neurotropism, and altered innate immune gene expression in mice. In contrast, vector competence studies showed no differences in transmission potential between ZIKV clones and unpassaged viruses.

## Results

### Mouse-adapted Zika viruses have increased transmission potential in *Aedes aegypti*

Previously (16), we performed in-vivo serial passage experiments in which a molecularly-barcoded ZIKV (ZIKV-BC), built on an Asian-lineage genetic backbone, was passaged in 5 parallel replicates (lineages) for 10 passages in *Ifnar1^-/-^* mice or in *Ae. aegypti* mosquitoes. To mimic natural host-cycling, ZIKV-BC was alternately-passaged in mice and mosquitoes for 10 passages in 5 parallel lineages. After subcutaneous inoculation of mice at passage 1, alternating passage was conducted via natural bloodfeeding transmission with small cohorts of mosquitoes feeding on an infected mouse, and then feeding on a naive mouse 12 days later (16). To evaluate the extent to which serial or alternate passage altered ZIKV replicative fitness and overall virulence, the phenotypes of the passaged viruses were evaluated in *Ifnar1^-/-^* mice and compared to unpassaged ZIKV-BC. Briefly, mouse-adapted viruses were universally lethal, whereas mosquito-adapted and alternate-passaged were heterogeneous in their outcomes (16). Next, to assess how serial and/or alternate passage affects vector competence for the adapted viruses, we compared the relative abilities of a subset of viruses from each passage series to be transmitted by *Ae. aegypti* in the laboratory. Two representative lineages from each passage series were chosen for vector competence experiments (see Table 1 in Methods): n=2 mouse-adapted ZIKVs, VP-A and VP-C; n=2 mosquito-adapted ZIKVs, MP-A and MP-E; and n=2 alternate-passaged ZIKVs, AP-A and AP-D. Since mouse passaged virus had uniform lethality, we chose two lineages with the most distinct frequencies of amino acid substitutions (VP-A: mixed population of NS2A-117T and −117V; VP-C: highest frequency of NS2A-117V and NS4A- E19G; see (16)). For the mosquito-adapted and alternate-passaged groups, we chose lineages that had distinct mortality phenotypes (see Table 1). To assess vector competence, mosquitoes were exposed to viremic bloodmeals (6-5-7.5 log_10_ PFU/mL) via water-jacketed membrane feeder maintained at 36.5 °C. Infection, dissemination, and transmission rates were assessed at 7, 14, and 21 days post feeding (dpf; n=39-80 per timepoint per virus group) using an in vitro transmission assay (16–18). Mosquitoes exposed to unpassaged ZIKV-BC served as experimental controls. Infection, dissemination, and transmission rates were similar for MP-A, MP-E, AP-A, AP-D, and ZIKV-BC at all timepoints. In contrast, VP-A and VP-C had significantly higher rates of infection at all time points compared to ZIKV-BC, with the exception of VP-A at 21d (two-tailed Fisher’s exact test) (Fig. 1A, green vs. black). Dissemination rates were also significantly increased across all timepoints for VP-A and VP-C compared to ZIKV-BC, with the exception of VP-A at 21d (Fig. 1B, green vs. black). Transmission rates of VP-C were significantly increased compared to ZIKV-BC at 21d (38% vs. 15%, p = 0.011, two-tailed Fisher’s exact test) (Fig. 1C). Thus, ZIKV serial passage in mice resulted in some variants displaying increased virulence coupled with increased transmission potential by mosquitoes.

**Figure 1.**
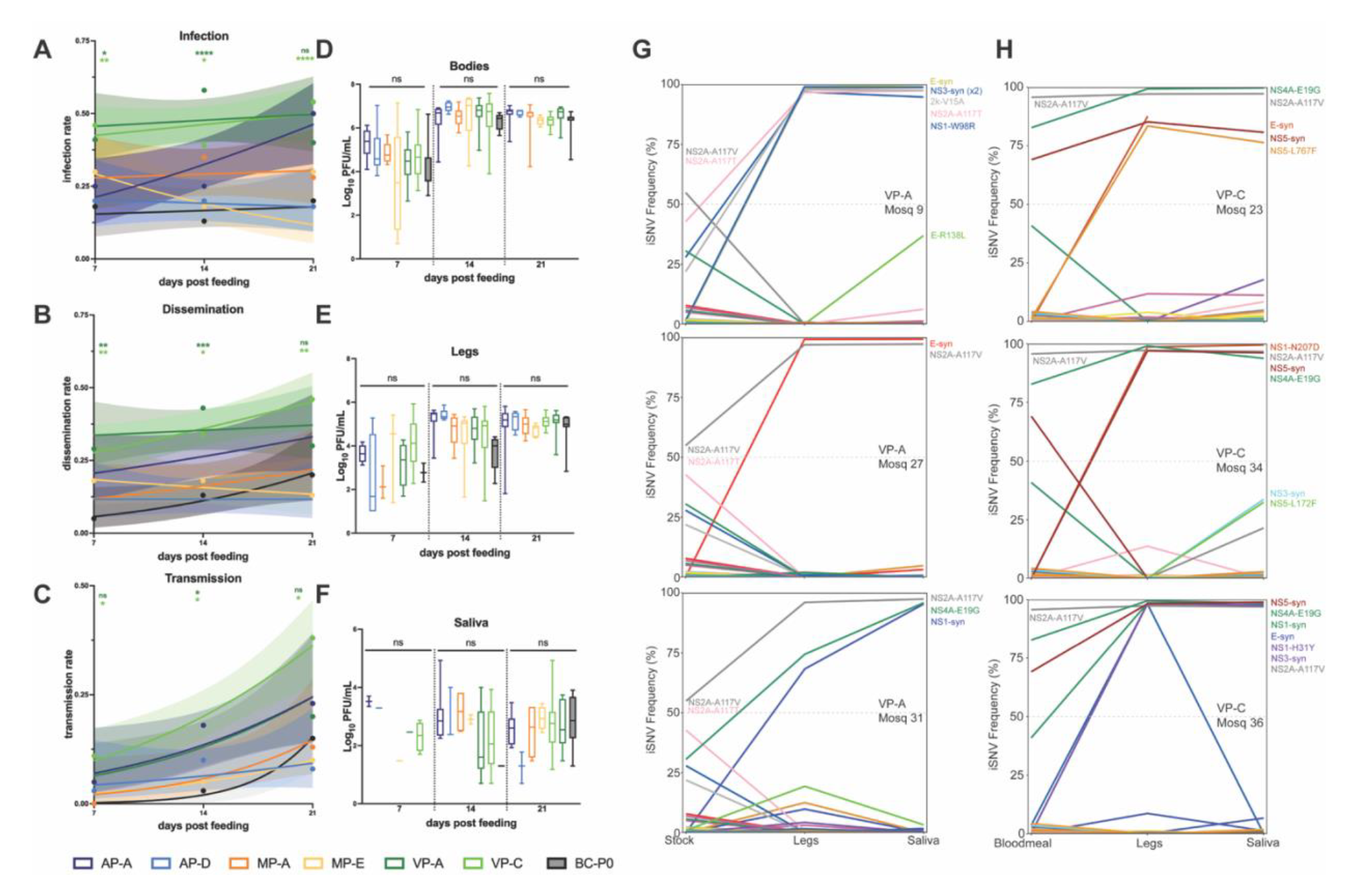
Vector competence of serially passaged Zika virus lineages. Female *Ae. aegypti* mosquitoes were exposed to passaged ZIKV strains via an artificial infectious bloodmeal and collected 7, 14, and 21 days post feeding (dpf). Infection **(a)** dissemination **(b)** and transmission **(c)** rates over the three collection time points are shown. Infection rate is the percentage of ZIKV-positive bodies, dissemination rate is the percentage of positive legs, and transmission rate is the percentage of positive saliva samples of total mosquitoes that took a bloodmeal (determined by plaque assay). Data points represent the empirically measured percentages (n=39-80 per data point). The lines represent the logistic regression results and the shaded areas represent the 95% confidence intervals of the logistic regression fits. Infection, dissemination, and transmission rates of VP-A and VP-C were compared to ZIKV-BC at each timepoint (two-tailed fisher’s exact test). *, *p* < 0.05; **, *p* < 0.01; ***, *p* < 0.001; ****, *p* < 0.0001; ns, not significant. Infectious virus was quantified via plaque assay from bodies **(d)** legs **(e)** and saliva **(f)** from all positive samples. Mean titers were not significantly different between virus groups in any tissue at any timepoint (one-way ANOVA with Tukey’s multiple comparisons test). ns, not significant. Paired legs and saliva samples from individual mosquitoes exposed to VP-A **(g)** and VP-C **(h)** 21 dpf were deep sequenced. Lines represent single nucleotide variant (SNV) frequency percentages between the stock or bloodmeal, legs, and saliva.

One possible mechanistic explanation for the shared fitness advantages in both mice and mosquitoes could be increased replicative capacity in both hosts. We therefore quantified infectious virus using plaque assays for all mosquito tissues that were positive in our vector competence assay and across all timepoints to assess whether the transmission advantage for VP-A and VP-C was due to increased replicative capacity of these viruses. We found no significant difference in infectious titers in any tissue between virus groups at any time point (one-way ANOVA with Tukey’s multiple comparisons test), and virus titers within virus groups were highly variable (Fig. 1D-F).

Next, we performed deep sequencing of virus populations replicating in legs (disseminated population) and saliva (transmitted population) from the VP-A- and VP-C-exposed mosquitoes at 21 dpf to confirm whether the amino acid substitutions selected for during mouse serial passage (NS2A-A117V, NS2A-A117T, NS2A-I139T, NS4A-E19G) were stably maintained in mosquito samples. A loss of these mutations would indicate that they are not advantageous in mosquitoes, and therefore not contributing to the enhanced transmission phenotype. The VP-A stock used to make infectious bloodmeals had a mixed population of V and T substitutions at NS2A position 117; in one mosquito (#9), NS2A-117T reached nearly 100% frequency in 21d saliva (Fig. 1G), whereas NSA-117V was present at nearly 100% frequency in the remaining saliva samples. In contrast, 2/3 VP-A mosquitoes lost NS4A E19G substitution while in 1/3, NS4A-19G was fixed in 21d saliva (Fig. 1G). Mosquitoes exposed to VP-C had nearly 100% NS2A-117V frequency in the stock and bloodmeal, and this substitution was maintained in all mosquitoes tested (Fig. 1H). VP-C mosquitoes also had NS4A-E19G present at nearly 100% frequency in the saliva at 21 dpf. NS2A-I139T, which was the lowest frequency variant, was not present in any mosquito samples sequenced.

### In vitro and in vivo characterization of mutant ZIKV clones

Phenotypic characterization of our experimentally evolved ZIKVs established that mouse-adapted viruses were more virulent in mice, more transmissible by mosquitoes, and that these viruses share amino acid substitutions that arose during serial passage in mice that are maintained during infection in mosquitoes. We therefore hypothesized that one, or a combination of, the amino acid substitutions NS2A-A117V, NS2A-A117T, or NS4A-E19G are contributing to the enhanced virulence and transmission phenotype. Since NS2A-I139T was the lowest frequency variant and was lost during mosquito infection, we did not include it in our panel. To assess the role of NS2A-A117V, NS2A-A117T, and NS4A-E19G in virulence and transmission, we engineered these substitutions, singly and in combination, into a Zika virus infectious clone derived from the Puerto Rican ZIKV isolate PRVABC59 (single mutants: NS2A- 117V, NS2A-117T, NS4A-E19G; double mutants: DM-E19G/A117V, DM-E19G/A117T). This is the same virus genetic backbone used in the previous experimental evolution study (16). Deep sequencing confirmed successful introductions of target mutations at a frequency >99%. Prior to use in vivo, we assessed viral infectivity and replication in vitro in Vero cells and C6/36 cells. All mutant viruses– as well as the control virus bearing the PRVABC59 isolate consensus sequence (ZIKV-IC)-- gave similar growth curves in both cell types (Fig. 2A-B). Of note, NS4A- E19G did reach higher significantly higher titers compared to ZIKV-IC at 48 and 72 hours post infection in C6/36 cells (hpi) (mixed-effects model with Holm-Sidak correction for multiple comparisons), but these differences were resolved by 96 hpi (Fig. 2B). Overall, these growth curves suggest that introducing these substitutions did not have a significant effect on either infectivity or replicative capacity in vitro.

**Figure 2:**
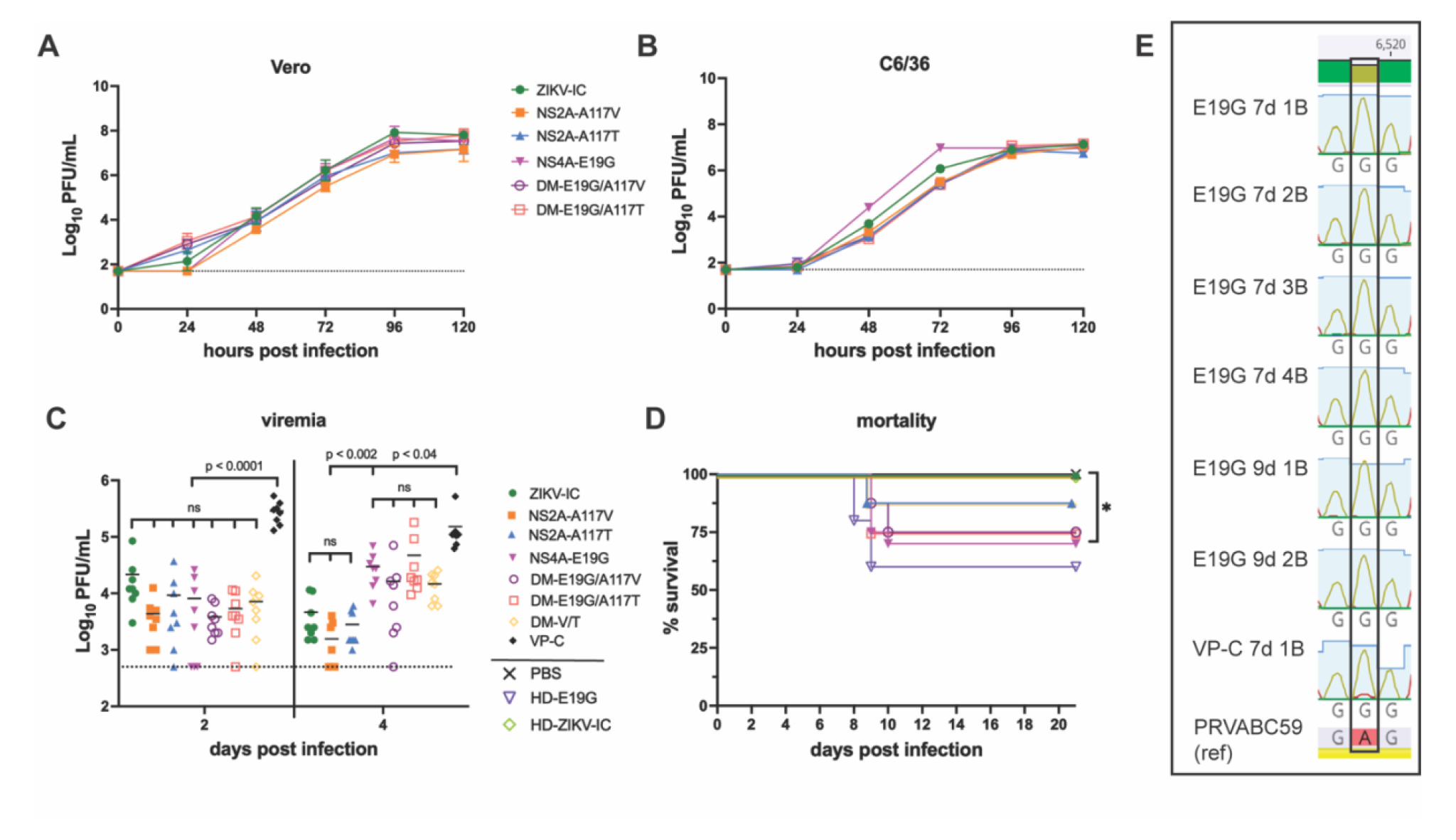
In vitro and in vivo characterization of ZIKV mutant clones. In vitro growth kinetics of ZIKV clones in Vero **(a)** and C6/36 **(b)** cells. Data points represent the mean of three replicates at each time point. Error bars represent standard deviation. **(c)** Serum viremia 2 and 4 days post infection (dpi) of *Ifnar1*^-/-^ mice inoculated with 10^3^ PFU of different ZIKV clones (n=8 for virus groups, n=4 for PBS control). Serum viremia from mutant clones was compared to previously published viremia data from VP-C infection. Differences in mean serum viremia between virus groups was compared by one-way ANOVA with Tukey’s multiple comparisons test. ns, not significant. The dotted line indicates the assay limit of detection. **(d)** Survival curves of *Ifnar1*^-/-^ mice inoculated with 10^3^ PFU of virus, or a PBS control. HD-E19G and HD-WTic groups were inoculated with 10^4^ PFU. Survival curves were compared to WTic by Fisher’s exact test. *, *p* < 0.05. **(e)** Chromatograms from Sanger sequencing of a subset of E19G 7d and 9d and VP-C 7d brains showing confirmation of the maintained NS4A position 19 substitution (nucleotide substitution at nt 6519 of the polyprotein: GAG → GGG). Sequenced amplicons were aligned to ZIKV-PRVABC59.

Next, to evaluate the extent to which the NS2A and NS4A amino acid substitutions alter the ZIKV virulence phenotype in vivo, we subcutaneously (s.c.) inoculated 5- to 7-week-old *Ifnar1^-/-^* mice in the footpad with 1×10^3^ PFU of the different ZIKVs (n=8-20), or PBS to serve as experimental controls (n=4). We also included a group inoculated with a 1:1 mixture of the double mutants (DM) to achieve a mixed population of NS2A-A117V and −A117T (DM-V/T) to assess whether diversity at this position was necessary for recapitulating the virulence phenotype. We collected serum 2 and 4 days post infection (dpi) to compare viremia kinetics between viruses. At 2 dpi, there were no significant differences in serum titers between the different ZIKVs (one-way ANOVA with Tukey’s multiple comparisons test) (Fig. 2C). At 4 dpi, serum titers did not differ significantly between ZIKV-IC, NS2A-A117V, and NS2A-A117T. However, NS4A-E19G and the double mutant viruses all reached significantly higher titers at 4 dpi compared to NS2A-A117V, NS2A-A117T, and ZIKV-IC (p < 0.002, one-way ANOVA with Tukey’s multiple comparisons test). Serum titers at 4 dpi in the NS4A-E19G and the double mutant infected groups did not differ significantly from each other (p > 0.13, one-way ANOVA). Additionally, when comparing these data to serum titers from mice infected with VP-C from (16), all mutant viruses and ZIKV-IC had significantly lower viremia at both timepoints (Fig. 2C).

Only the NS4A-E19G-inoculated groups showed significant mortality compared to ZIKV-IC and PBS-inoculated, age-matched controls (30% mortality, log-rank test, *p* = 0.02) (Fig. 2D). DM- E19G/A117V- and DM-E19G/A117T-groups showed similar rates of mortality (25%). Because NS4A-E19G did not recapitulate the 100% mortality phenotype observed with the uncloned mouse-adapted viruses (16), we inoculated a group of mice (n=5) with a higher challenge dose (10^4^ PFU) of NS4A-E19G or ZIKV-IC. The higher-dose, NS4A-E19G-inoculated group resulted in a 40% mortality rate (Fig. 2D). Synthesizing the mortality data from all infections with viruses containing NS4A-E19G (low-dose and high-dose NS4A-E19G and double mutants) compared to those without this substitution (ZIKV-IC and NS2A single mutants), we conclude that the virulence phenotype is NS4A-E19G-dependent.

Given the incomplete penetrance of the NS4A-E19G phenotype, we posited that mice that this substitution could have at least partially reverted to wild type in mice that did not succumb to infection and had lower levels of virus detectable in the brain. We therefore sequenced virus from brain tissue of mice infected with NS4A-E19G 7 and 9-10 dpi and determined that this substitution was stably maintained during mouse infection (Fig. 2E).

### Increased neurotropism and innate immune gene expression is NS4A-E19G-dependent

ZIKV NS4A specifically has been shown to antagonize interferon induction via interactions with the RIG-I-like-receptors (RLRs), retinoic acid-inducible gene I (RIG-I), and melanoma-differentiation-associated gene 5 (MDA-5), which are cytosolic RNA sensors that sense viral RNAs during infection and trigger signaling pathways resulting in the production of type I interferon and proinflammatory cytokines (19, 20). Since our experiments were done in mice that lack type I IFN receptor 1 function, we expect that this antagonism occurs via RLR signaling through the mitochondrial antiviral-signaling protein (MAVS) pathway (21). Indeed, increasing evidence suggests that viruses can escape the host antiviral response by interfering at multiple points in the MAVS signaling pathway, thereby maintaining viral replication and expanding tropism (22). MAVS signaling is initiated by RLRs that recognize viral RNA, therefore we posit that NS4A-E19G confers an enhanced ability to evade innate immune responses in mice resulting in higher titers and broader tissue tropism, including neuroinvasion (23–25).

To evaluate whether NS4A-E19G expands virus tropism, we s.c. inoculated *Ifnar1^-/-^*mice with 10^3^ PFU of NS4A-E19G, ZIKV-IC, or the mouse-adapted virus lineage, VP-C. Brains were collected at 3 or 7 dpi, and from animals that met endpoint criteria (∼9-10 dpi) from experiments described above (Fig. 2D). Brains were also collected at 9 dpi from a group of ZIKV-IC- inoculated mice to serve as controls for the 9-10 dpi timepoint. Brains from VP-C-inoculated animals could not be collected from later timepoints because all mice met endpoint criteria by 7 dpi (see (16)). Infectious virus was quantified from brain homogenate via plaque assay, and viral RNA (vRNA) loads were evaluated via quantitative reverse transcription PCR (qRT-PCR). Viral loads in the brain were significantly higher in mice infected with NS4A-E19G compared to ZIKV- IC at 3dpi (*p* = 0.001, one-way ANOVA with Tukey’s multiple comparisons test), while mean viral load did not differ between NS4A-E19G and VP-C at 3 dpi (Fig. 3A). Viral load increased in all groups from 3 to 7dpi, with no significant difference in brain vRNA loads between ZIKV-IC and NS4A-E19G-inoculated groups (Fig. 3A). At 9-10 dpi, vRNA loads continued to increase in animals infected with NS4A-E19G or double mutants, but did not differ significantly from VP-C- inoculated groups at 7 dpi. ZIKV-IC viral loads were significantly lower than NS4A-E19G and double mutants at 9 dpi (*p* < 0.001). Similar patterns were observed in infectious virus titers in the brain (Fig. 3B). Infectious virus was only detectable at low levels in 2 ZIKV-IC-infected brains, 2 samples at 3 dpi and 4 samples at 7 dpi from NS4A-E19G-inoculated groups. Although not statistically different from ZIKV-IC at 7dpi, the number of plaque-assay-positive brains from NS4A-inoculated animals does roughly parallel the mortality rate observed in this group. At 9-10 dpi, all NS4A-E19G and double mutant infected groups had detectable infectious virus in the brain, while no ZIKV-IC brains were positive (Fig. 3B). While discrepancies in patterns between vRNA and infectious titers are likely due to factors such as differential assay sensitivity and qRT-PCR detecting partial and non-viable RNA fragments, both assays show increased neurotropism of viruses possessing NS4A-E19G.

**Figure 3:**
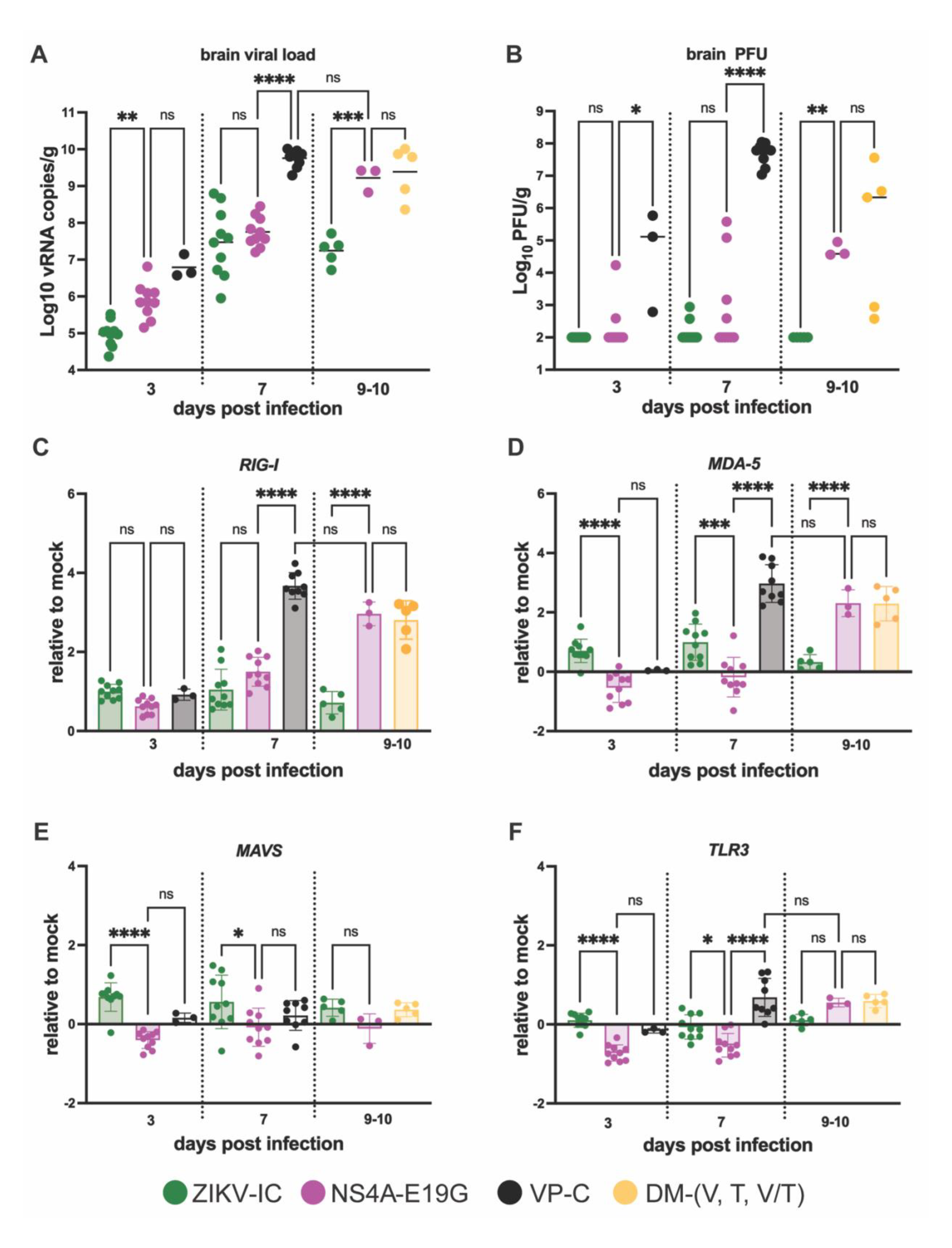
Differential neurotropism and innate immune gene responses to ZIKV mutants. Viral RNA **(a)** and infectious virus **(b)** were quantified from brain tissue collected 3, 7, and 9-10 days after 10^3^ PFU inoculation of *Ifnar1^-/-^* mice with ZIKV-IC, NS4A-E19G, VP-C or, double mutant clones by qRT-PCR and plaque assay respectively. Transcript abundance of *RIG-I* **(c),** *MDA-5* **(d)**, *MAVS* **(e)**, and *TLR3* **(f)** was analyzed from brains collected 3, 7, and 9-10 dpi by qPCR. Expression levels were normalized to ActB and Hprt and the ddC_T_ value was calculated as log2 fold change expression relative to mock-inoculated mice. Data points represent individual samples. Means with standard deviation are shown. Viral loads, infectious viral titers, and relative fold change of transcript abundance were compared between virus groups at each timepoint by one-way ANOVA with Tukey’s multiple comparisons test. *, *p* < 0.05; **, *p* < 0.01; ***, *p* < 0.001; ****, *p* < 0.0001; ns, not significant.

Next, to evaluate whether NS4A-E19G modulates antiviral signaling in the brain, we measured the relative transcript abundance of *RIG-I*, *MDA-5*, *MAVS*, and Toll-like receptor 3 (*TLR3*) in the brain at 3, 7, and 9-10 dpi. *TLR3* was included as an additional pattern recognition receptor (PRR), because it detects double-stranded RNA and has been shown to modulate the inflammatory response to ZIKV infection in astrocytes, as well as interact with the RIG-I pathway (26, 27). Additionally, in the context of other arboviruses, TLR3 has been shown to be both protective and pathologic during West Nile virus (WNV) infections in the brain (28, 29).

At 3 dpi, we observed modest induction of *RIG-I* for all virus groups compared to mock-inoculated controls (Fig. 3C). At 7 dpi, ZIKV-IC and NS4A-E19G had similar *RIG-I* expression profiles that were significantly lower than VP-C. However, at 9-10 dpi, viruses containing NS4A- E19G had significantly higher *RIG-I* transcript abundance compared to ZIKV-IC (*p* < 0.0001, one-way ANOVA with Tukey’s multiple comparisons test). *MDA-5* expression patterns were similar to *RIG-I*, with increased *MDA-5* abundance detected at 7 dpi in VP-C and at 9-10 dpi for viruses containing NS4A-E19G (Fig. 3D). Interestingly, at 3 and 7 dpi, brains of mice infected with NS4A-E19G had significantly less *MDA-5* transcript abundance compared to those of mice infected with ZIKV-IC (*p* < 0.0002, one-way ANOVA with Tukey’s multiple comparisons). The *MDA-5* expression pattern suggests that the NS4A-E19G substitution may contribute to more efficient suppression of *MDA-5* signaling early in infection. Alternatively, increased abundance of both *RIG-I* and *MDA-5* at later timepoints in brains from both VP-C and NS4A-E19G groups suggests that prolonged innate immune activation may be responsible for adverse outcomes in these mice. In contrast, *MAVS* and *TLR3* were not induced at the later timepoints for any virus group, but brains of mice infected with NS4A-E19G had significantly less *MAVS* and *TLR3* transcript abundance at 3 and 7 dpi compared to ZIKV-IC (*p* < 0.0001, one-way ANOVA with Tukey’s multiple comparisons test) (Fig. 3E-F). Early suppression of *MDA-5*, *MAVS*, and *TLR3* suggests that NS4A-E19G may contribute to an enhanced ability to evade early innate immune responses, and thus broader tissue tropism, which we observed as increased neuroinvasion, mortality, and viral replication with viruses containing NS4A-E19G. Alternatively, significant induction of *RIG-I* and *MDA-5* later in infection indicates prolonged activation of RLR signaling in the brain, which may contribute to the virulence phenotype for NS4A-E19G viruses (29, 30). Although the exact mechanism cannot be resolved from these data, the results clearly show differences in the modulation of innate immune signaling for viruses containing NS4A-E19G compared to ZIKV-IC.

### Transmission potential is not impacted by NS2A-117V, NS2A-117T, or NS4A-19G

Finally, to evaluate the extent to which the NS2A and NS4A amino acid substitutions altered the vector competence phenotype, we compared the relative abilities of NS4A-E19G, DM- E19G/A11V, DM-E19G/A117T, and ZIKV-IC to be transmitted by *Ae. aegypti*.

We focused on viruses containing the NS4A-E19G substitution because of its role in the virulence phenotype in mice (Fig. 2). Because vector competence for the double mutants did not significantly differ from the ZIKV-IC control, the single NS2A mutants were not pursued in vector competence experiments. Vector competence was performed as described above. Briefly, female *Ae. aegypti* mosquitoes were exposed to bloodmeals containing either ZIKV-IC, NS4A- E19G, DM-E19G/A11V, DM-E19G/A117T, as well as a bloodmeal that contained a 1:1 mixture of the two double mutants (DM-V/T). Bloodmeal titers ranged from (6.6-7.3 log_10_ PFU/mL). Infection, dissemination, and transmission rates were assessed at 7, 14, and 21 dpf (n=20-80 per timepoint per virus group). ZIKV-IC and NS4A-E19G exposures were repeated in two independent replicates with distinct generations of mosquitoes to account for stochastic variation between generations. No differences were observed in infection, dissemination, or transmission rates at any timepoint between viruses (Fig. 4A-C).

**Figure 4:**
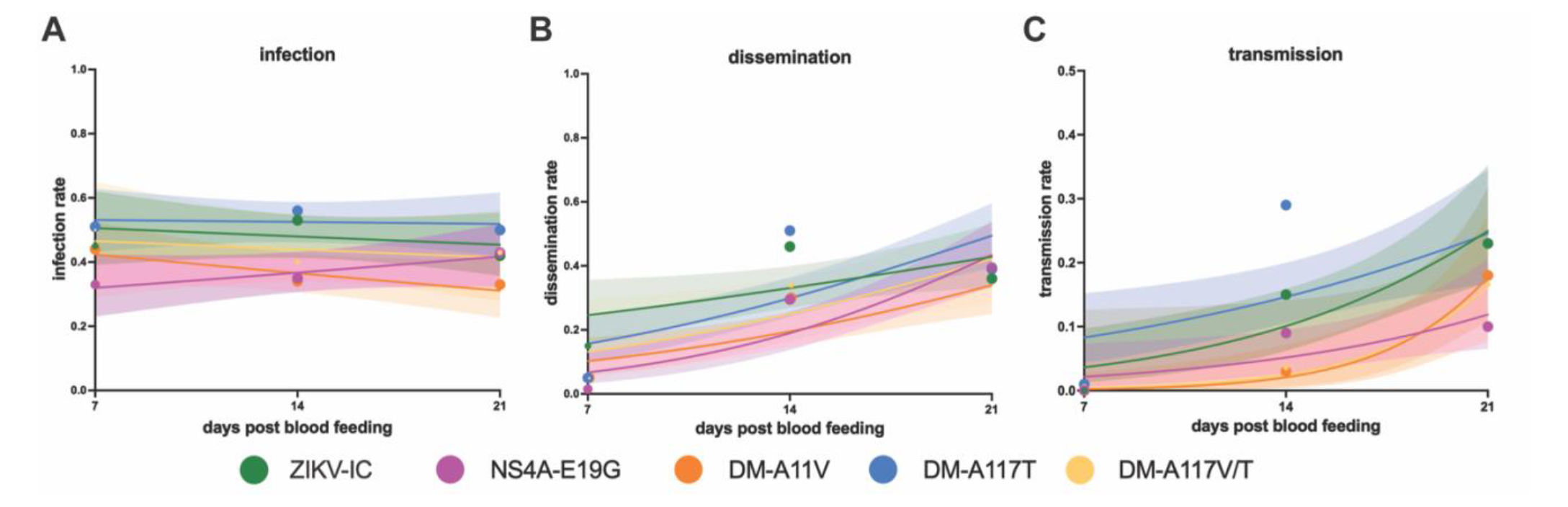
Vector competence of ZIKV mutants. Female *Ae. aegypti* mosquitoes were exposed to passaged ZIKV strains via an artificial infectious bloodmeal and collected 7, 14, and 21 days post feeding (dpf). Infection **(a)** dissemination **(b)** and transmission **(c)** rates over the three collection time points are shown. Infection rate is the percentage of ZIKV-positive bodies, dissemination rate is the percentage of positive legs, and transmission rate is the percentage of positive saliva samples (determined by plaque assay screen). Data points represent the empirically measured percentages (n= 20-80 per data point, point size is proportional to group size). The lines represent the logistic regression results and the shaded areas represent the 95% confidence intervals of the logistic regression fits.

## Discussion

Here, we expanded on our previous work (16), to demonstrate that ZIKV adaptation to mice, which increased virulence, is coupled with increased transmission potential by mosquitoes. These observations are important because knowledge on how flaviviruses limit or avoid the effects of trade-offs from dual-host cycling is limited, and thus providing further evidence that flaviviruses are capable of limiting the effects of trade-offs from host cycling. This finding is consistent with observations from other studies that mostly refute the trade-off hypothesis for arboviruses (3, 4, 31, 32). For example, serial passaging of WNV in chicks resulted in avian-passaged WNV acquiring fitness gains in both chicks and *Culex pipiens* mosquitoes (31). Similarly, experiments with single-host passaged St. Louis encephalitis virus found that some avian-passaged SLEVs displayed fitness gains in chicks without fitness losses in mosquitoes (32). We also found that during direct vertebrate transmission chains, ZIKV consistently evolved these enhanced virulence and transmission properties coincident with the selection of two polymorphisms at a single site in NS2A—A117V and A117T—and at NS4A E19G (16). NS2A 117V is present on naturally isolated ZIKV genomes from humans and *Ae. aegypti* reported in the Bacterial and Viral Bioinformatics Resource Center (BV-BRC), indicating it is a viable variant in natural transmission cycles. Importantly, NS4A E19G has also been identified in a naturally isolated ZIKV genome from a pregnant woman in Honduras in 2016, reported in Nextstrain and has not been characterized prior to this study. In contrast, the NS2A 117T variant has not been previously detected in natural or laboratory isolates to our knowledge (BV-BRC database: https://www.bv-brc.org; Nextstrain: https://nextstrain.org). The emergence of naturally occurring variants within our passaging experiments supports the biological relevance of this experimental system to mimic natural transmission dynamics and produce variants viable in nature which warrant further characterization.

All studies seeking to link viral genotype to phenotype using reverse genetics can be limited by the genetic context of the cloned virus(es) chosen for analysis. As a result, it may be that these substitutions examined in the context of different ZIKV strains might have different phenotypic effects. The parental virus used in our studies was PRVABC59, a 2016 isolate from Puerto Rico. A previous study found that NS2A A117V enhanced virulence of two additional strains of American-sublineage ZIKV in mice (33). However, we did not observe a significant difference in pathogenic potential of ZIKV encoding alanine or valine at NS2A position 117. Flavivirus NS2A and NS4A have been previously shown to act as interferon (IFN) antagonists (34–36) and as a result are mediators of flavivirus pathogenesis (37, 38) the virulence phenotype was NS4A- E19G-dependent.

Flavivirus NS4A is a multifunctional protein involved in membrane remodeling and the establishment of the viral replication complex (39, 40), innate immune antagonism (19), as well as modulating autophagy, which is important in neurodevelopment (41, 42). Position 19 falls within the cytoplasmic N terminus of NS4A, a region particularly important for protein stability and homo-oligomerization (43). While studies of ZIKV NS4A are limited, studies investigating the role of this region of NS4A in both DENV and WNV found that mutations introduced into this region were highly attenuating to viral replication (44–46). Another study, using a yellow fever virus strain with defective replicative capacity due to a large in-frame deletion in the NS1 gene, found that replication could be restored by an adaptive mutation in the N-terminal region of NS4A (47). Previous studies identified interactions between NS4A and NS2A which promote neurovirulence and innate immune antagonism (19). We had therefore posited that there may be an epistatic interaction between the NS2A and NS4A substitutions. Our results, however, show similar outcomes from clones containing NS4A-E19G with or without NS2A substitutions, suggesting there is no epistatic interaction between the NS4A E19G and NS2A substitutions— at least in the environments tested here— nor do these mutations have functionally redundant roles in promoting neurovirulence.

Little is known about which regions of the protein are involved in NS4A-mediated immune regulation. We identified distinct patterns of innate immune gene expression in the brain between ZIKV-IC-infected brains and NS4A-E19G-infected brains. We therefore speculate that increased and prolonged immune activation in the brain of NS4A-E19G-infected mice contributes to immunopathology due to increased proinflammatory cytokine production from RLR signaling (48–50). Numerous proteins—such as a subset of TRIM proteins—negatively regulate the RIG-I pathway to protect against prolonged signaling-induced immunopathology, so it is possible that the NS4A substitution facilitates improved antagonism of negative regulatory proteins of these immune pathways (51, 52). Our data suggest that increased replication and neurotropism likely contribute to this change in innate immune signaling, but more studies are needed to disentangle the exact underlying mechanism(s). We do, however, provide further support for the multifaceted role of NS4A in both replication and immune modulation.

NS4A may also play a functionally redundant and evolutionarily conserved role in the mosquito host. There are considerable similarities in insect and mammalian cytosolic sensing of RNA; RIG-I-like antiviral proteins have been identified in flies, including mosquitoes (53–56). In mammals, RLRs are critical for the production of type I interferon in cells; whereas, in flies Dicer-2 appears to be a functional equivalent to mammalian RLRs (53). The mammalian RLRs retinoic RIG-I and MDA-5 sense cytosolic viral RNAs during infection and activate signaling pathways that culminate in the production of type I IFNs and proinflammatory cytokines as part of the antiviral immune response (57, 58). In mosquitoes, Vago appears to function as an IFN-like antiviral cytokine that is activated in a Dicer-2 dependent manner (54). However, since neither polymorphism in NS2A or NS4A impacted mosquito vector competence (Fig. 4), we do not have robust evidence for this in our system.

Importantly, the virulence phenotype we observed with NS4A-E19G was not 100% penetrant. Previously, it has been shown that viral population diversity is important for fitness, virulence, and pathogenesis (59–61). For example, polio viruses with reduced viral diversity had an attenuated phenotype in mice and also lost neurotropism (59). However, when we mixed both double mutants (DM-E19G/A117V and DM-E19G/A117T) for mosquito vector competence and mouse pathogenesis experiments, the mixed populations had no impact on the mosquito vector competence phenotype and were similarly pathogenic to the single NS4A-E19G virus. Although both passaged and clone viruses were constructed on the same parental backbone (ZIKV- PRVABC59), the passaged-viruses contained a molecular barcode and thus barcode diversity while the mutant ZIKVs used here did not. While we would not expect silent barcodes to enhance virulence, and there is no evidence for this when comparing the virulence phenotype of the barcoded virus versus the ZIKV-IC used here (16, 62), the accumulation of genetic diversity may be important for the virulence phenotype. Critically, we were not able to investigate all low-frequency variants present in the mouse-passaged viruses. For example, NS2A I139T was found in all five lineages but was only found on less than 50% of the ZIKV genomes (16). We chose not to investigate this variant because NS2A I139T was not detected when we put VP-A and VP-C into mosquitoes (Fig. 1H).

Since the NS4A-E19G virus was unable to completely recapitulate the virulence phenotype observed with the mouse-adapted viruses from our previous study (16), we posit that this phenotype is multifactorial, involving some combination of the E19G amino acid substitution, virus population diversity and codon usage—all of which are supported by the work presented herein and by the analyses presented in our previous paper (16). Additionally, like most studies, we focused on the coding region of the genome. The untranslated 5’ and 3’ ends of the flavivirus genome (5’ and 3’ UTRs), however, have gained increased focus for the multifaceted roles of UTR-derived subgenomic flavivirus RNAs (sfRNA) in viral replication, mosquito transmission, as well as viral pathogenicity in vertebrate and invertebrate hosts (63–66). In fact, a 2015 study identified 2 substitutions within the 3’ UTR of an emergent strain of DENV, which resulted in increased production of sfRNAs that reduced RIG-I signaling and subsequent IFN induction, and ultimately contributed to greater epidemiological fitness of this DENV strain (67). Future study of the evolutionary trajectory of ZIKV, as well as other RNA viruses, should consider these alternative genetic components as potential determinants for changes in viral phenotype. The future spread of ZIKV is unpredictable, however, these studies collectively show that direct transmission chains could enable the emergence of more virulent and transmissible ZIKV strains. While the underlying genetics of these adaptations remain unclear, these studies highlight the capacity of flaviviruses to adapt in ways which limit the restrictions of dual host cycling.

## Materials and Methods

### Ethical approval

This study was approved by the University of Minnesota, Twin Cities Institutional Animal Care and Use Committee (Protocol Number, 1804–35828).

### Cells and viruses

African Green Monkey kidney cells (Vero; ATCC #CCL-81) were maintained in Dulbecco’s modified Eagle medium (DMEM) supplemented with 10% fetal bovine serum (FBS; Hyclone, Logan, UT), 2 mM L-glutamine, 1.5 g/L sodium bicarbonate, 100 U/ml penicillin, 100 μg/ml of streptomycin, and incubated at 37 °C in 5% CO2. *Aedes albopictus* mosquito cells (C6/36; ATCC #CRL-1660) were maintained in DMEM supplemented with 10% fetal bovine serum (FBS; Hyclone, Logan, UT), 2 mM L-glutamine, 1.5 g/L sodium bicarbonate, 100 U/ml penicillin, 100 μg/ml of streptomycin, and incubated at 28 °C in 5% CO2. Cell lines were obtained from the American Type Culture Collection and were tested and confirmed negative for mycoplasma. Viruses were generated by in vivo passaging of a barcoded Zika virus infectious clone derived from the PRVABC59 strain followed by cell culture amplification after 10 passages as previously described (16). For vector competence experiments with passaged viruses, amplified passage 10 viruses were passed through another round of cell culture amplification and then concentrated using Amicon Ultra-15 Centrifugal filter unit (Millipore Sigma) to achieve necessary infectious blood meal titers. Strains used for these experiments are presented in Table 1 (below).

**Table 1.** Viruses used for vector competence (Fig. 1).

| **Virus** | **Passage group** | **Mortality in mice (%)<sup>\*</sup>** |
| --- | --- | --- |
| AP-A | Alternate | 100 |
| AP-D | Alternate | 0 |
| MP-A | Mosquito | 75 |
| MP-E | Mosquito | 0 |
| VP-A | Mouse | 100 |
| VP-C | Mouse | 100 |
| ZIKV-BC | Unpassaged | 12.5 |

<sup>\*</sup>Mortality determined in prior study, Riemersma et al. 2020 (14).

### Generation of ZIKV mutant clones

Mutagenic primers were designed using SnapGene 6.0.2 software (GSL Biotech). PCRs were performed using the SuperFi II Master Mix (Thermo Fisher 12368010), and the products were gel purified using the Machary-Nagel NucleoSpin Gel and PCR Clean-up kit (740609). Amplicons were assembled using the NEB Builder HiFi DNA Assembly Master Mix (E2621L). The assembly products were digested with DpnI (R0176S), Lambda exonuclease (M0262S), and Exonuclease I (M0293S) and amplified using SuperPhi RCA Premix Kit with Random Primers (Evomic Science catalog number PM100) (68). RCA products were then transfected into HEK293A cells (69). The supernatant was harvested, clarified by centrifugation, and sequenced by Sanger sequencing to confirm the mutation of interest was introduced.

### Viral sequencing

#### Whole genome sequencing (WGS) and analysis

ZIKV mutant clones were additionally deep-sequenced to further confirm that no other polymorphisms were detected in these stocks, as previously described (70).

For sequencing from mosquito tissue samples, viral RNA was extracted from 50uL of saliva and 200uL of bodies, virus stocks, and blood meals. All RNA extractions were performed on a Maxwell 16 instrument (Promega) with the Maxwell RSC Viral Total Nucleic Acid Purification Kit and eluted in 50uL of nuclease-free water. Whole genome ZIKV sequencing libraries were then prepared and sequenced as previously described (16). WGS libraries were generated using the previously described tiled PCR amplion approach (71, 72). WGS libraries were sequenced with paired-end 150-bp reads on an Illumina NovaSeq 6000 by the UW Biotechnology Center. Similarly, WGS analysis for mosquito samples was conducted as previously described (16). Briefly, raw reads were quality-trimmed, merged, and aligned to the ZIKV PRVABC59 reference strain. Consensus sequences were extracted and variants were called against reference and consensus sequences. Frequency of single nucleotide variants (SNVs) across paired mosquito samples were analyzed using previously developed pipelines. Data processing, analysis, and visualization scripts are publicly available (https://github.com/tcflab/ZIKVBC_HostCycling).

#### Sanger sequencing

A 388 bp region surrounding NS4A position 19 was amplified from total RNA extracted from a subset of brains using OneStep RT-PCR (Qiagen) using the following primers: Forward: (5’- AGAGTGCTCAAACCGAGGTG-3’), Reverse: (5’- TTTCCGAGAGCCACATGAGC-3’). Amplicons were sequenced directly by Sanger sequencing (Azenta Life Sciences). Sequences were viewed with Geneious Prime.

### Plaque assay

ZIKV screens from mouse serum and tissue, mosquito tissue, in vitro growth curves, and titrations for virus quantification from virus stocks were completed by plaque assay on Vero cell cultures. Duplicate wells were infected with 0.1 ml aliquots from serial 10-fold dilutions in growth media and virus was adsorbed for 1 h. Following incubation, the inoculum was removed, and monolayers were overlaid with 3 ml containing a 1:1 mixture of 1.2% oxoid agar and 2X DMEM (Gibco, Carlsbad, CA) with 10% (vol/vol) FBS and 2% (vol/vol) penicillin/streptomycin. Cells were incubated at 37°C in 5% CO2 for four days for plaque development. Cell monolayers then were stained with 3 ml of overlay containing a 1:1 mixture of 1.2% oxoid agar and 2X DMEM with 2% (vol/vol) FBS, 2% (vol/vol) penicillin/streptomycin, and 0.33% neutral red (Gibco). Cells were incubated overnight at 37 °C and plaques were counted.

### Virus isolation

Total RNA was extracted and purified from homogenized brain tissue using a Direct-zol RNA kit (Zymo Research) and used for onward analysis of vRNA quantity and gene expression. RNA was then quantified using quantitative RT-PCR.

### Quantitative Reverse Transcription PCR (qRT-PCR)

Viral RNA was quantified from extracted total RNA from brain tissue by quantitative reverse transcription-PCR (RT-qPCR) as described previously (10). The RT-PCR was performed using the Taqman Fast Virus 1-step master mix (Applied Biosystems, Foster City, CA) on a QuantStudio3 (ThermoFisher). The viral RNA concentration was determined by interpolation onto an internal standard curve composed of seven 10-fold serial dilutions of a synthetic ZIKV RNA fragment. The ZIKV RNA fragment is based on a ZIKV strain derived from French Polynesia that shares >99% identity at the nucleotide level with the Puerto Rican lineage strain from which all viruses were derived that are used in the infections described in this report.

### Gene expression

cDNA was synthesized using the High-Capacity RNA-to-cDNA kit (Applied Biosystems) from RNA extracted from brain homogenate as previously described. Gene expression of RIG-I, MDA-5, MAVS, and TLR3 was then quantified from cDNA by qPCR using customized TaqMan® Array Mouse Antiviral Response 96-well plates (ThermoFisher) run on a QuantStudio3 (ThermoFisher). Innate immune gene transcript abundance was normalized to *ActB and Hprt* and then the threshold cycle value (2-delta delta *CT*) was calculated relative to mock-inoculated controls.

### In vitro viral replication

Six-well plates containing confluent monolayers of Vero and C6/36 cells were inoculated with each clone virus (WTic, NS4A-E19G, NS2A-A117V, NS2A-A117T, DM-E19G/A117V, DM-E19G/A117T) in triplicate at a multiplicity of infection of 0.01 PFU/cell. After one hour of adsorption at 37°C, inoculum was removed and the cultures were washed three times. Fresh media were added and the cell cultures were incubated for 5 days at 37°C with aliquots removed every 24 hours and stored at −80°C. Viral titers at each time point were determined by plaque assay on Vero cells.

### Mice

*Ifnar1^−/−^* mice on the C57BL/6 background were bred in the specific pathogen-free animal facilities of the University of Minnesota College of Veterinary Medicine. Mixed sex groups of five to seven week old mice were used for all experiments.

#### Subcutaneous inoculation

*Ifnar1^−/−^* mice were subcutaneously inoculated in the left hind footpad with 10^3^ PFU of the selected ZIKV clone, or VP-C in 20ul of sterile phosphate buffered saline (PBS) or with sterile PBS alone as experimental controls. An inoculum titer or 10^3^ PFU was selected to match the prior study of passaged ZIKV viruses in mice (CITE). An additional 10^4^ PFU inoculation group was included for WTic and NS4A-E19G. Submandibular blood draws were collected 2 and 4 dpi to confirm viremia and compare replication kinetics. Mice were weighed daily and monitored for clinical signs of morbidity. Mice were euthanized at the experimental endpoint of 21 dpi, or if they met early endpoint euthanasia criteria.

#### Mouse necropsy

Following inoculation with WTic, NS4A-E19G, or VP-C, mice were sacrificed at 3 and 7 dpi, or if they met early endpoint criteria. Immediately following euthanasia, brain tissue was carefully dissected using sterile instruments, placed in a culture dish, rinsed with sterile PBS, and then brains were individually collected in pre-weighed tubes of PBS supplemented with 20% FBS and penicillin/streptomycin. These tissues were then weighed and homogenized by either Omni homogenizer (Omni International, Omni Tissue Homogenizer (TH) 115V) or by 5-mm stainless steel beads with a TissueLyser (Qiagen) and used for onward analysis of infectious virus and vRNA quantification, and gene expression analysis.

### Mosquito strain and colony maintenance

All mosquitoes used in this study were maintained at the University of Minnesota, Twin Cities as previously described (73) in an environmental chamber at 26.5 ± 1 °C, 75% ± 5% relative humidity, and with a 12 h light and 12 h dark photoperiod with a 30 min crepuscular period at the beginning of each light cycle. The *Aedes aegypti* Gainesville strain used in this study was obtained from Benzon Research (Carlisle, PA), and was originally derived from the USDA “Gainesville” strain. Three-to six-day-old female mosquitoes were used for all experiments.

### Vector competence studies

Mosquitoes were exposed to virus-infected bloodmeals via water-jacketed membrane feeder maintained at 36.5 °C (74). Bloodmeals consisted of defibrinated sheep blood (HemoStat Laboratories, Inc.) and virus stock, yielding infectious bloodmeal titers ranging from (insert) PFU/ml. Bloodmeal titer was determined after feeding. Infection, dissemination, and transmission rates were determined for individual mosquitoes and sample sizes were chosen using well-established protocols (17, 18, 75, 76). Briefly, mosquitoes were sucrose starved overnight prior to bloodmeal exposure. Mosquitoes that fed to repletion were randomized, separated into cartons in groups of 40–50, and maintained on 0.3 M sucrose in a Conviron A1000 environmental chamber at 26.5 ± 1 °C, 75% ± 5% relative humidity, with a 12 h photoperiod within the Veterinary Isolation Facility BSL3 Insectary at the University of Minnesota, Twin Cities. All samples were screened by plaque assay on Vero cells.

### Statistical analyses

All statistical analyses were conducted using GraphPad Prism 9 (GraphPad Software, CA, USA). Fisher’s exact test was used to determine significant differences in vector competence between viruses in each tissue at each timepoint. For survival analysis, Kaplan-Meier survival curves were analyzed by the log-rank test. One-way ANOVA with Tukey’s multiple comparisons was used to determine differences in viremia, brain viral loads and titers, and relative transcript abundance between virus groups.

## Data availability

Raw Illumina sequencing data will be available on the NCBI Sequence Read Archive, BioProject number pending.

## Acknowledgements

We acknowledge the University of Minnesota, Twin Cities BSL3 Program for facilities and Neal Heuss for support. This work was supported by NIH grants R21AI131454 and R01AI132563. A.S.J. was supported by T32 AI083196 from the National Institute of Allergy and Infectious Disease. The Wisconsin National Primate Research Center is supported by grants P51RR000167 and P51OD011106.

